# TMIGD1, a putative tumor suppressor, induces G2-M cell cycle checkpoint arrest in colon cancer cells

**DOI:** 10.1101/2020.06.06.138057

**Authors:** Kyle Oliver Corcino De La Cena, Rachel Xi-Yeen Ho, Razie Amraei, Nick Woolf, Joseph Y. Tashjian, Qing Zhao, Sean Richards, Josh Walker, Juanni Huang, Vipul Chitalia, Nader Rahimi

## Abstract

Colorectal cancer (CRC) is a leading non-familial cause of cancer mortality among men and women. Although various genetic and epigenetic mechanisms have been identified, the full molecular mechanisms deriving CRC tumorigenesis remains incompletely understood. In this study, we demonstrate that cell adhesion molecule transmembrane and immunoglobulin domain containing1 (TMIGD1) is highly expressed in mouse and human normal intestinal epithelial cells. We have developed TMIGD1 knockout mice and show that the loss of TMIGD1 in mice results in the development of adenomas in small intestine and colon. Additionally, the loss of TMIGD1 in mouse impaired intestinal epithelium brush border formation, junctional polarity and maturation. Mechanistically, TMIGD1 inhibits tumor cell proliferation, cell migration, arrests cell cycle at G2/M phase and induces expression of p^21CIP1^ (cyclin-dependent kinase inhibitor 1), and p^27KIP1^ (cyclin-dependent kinase inhibitor 1B) expression, key cell cycle inhibitor proteins involved in the regulation of the cell cycle. Moreover, we demonstrate that TMIGD1 is progressively downregulated in sporadic human CRC and correlates with poor overall survival. Our findings identify TMIGD1 as a novel tumor suppressor gene and provide insights into the pathogenesis of colorectal cancer and possibilities as a potential therapeutic target.

## Introduction

Colorectal cancer (CRC) is the second most common malignancy in western countries^1^. CRC’s high mortality is associated with tumor metastasis and poor response of patients to current standard-of-care drugs^2-4^. The development of CRC is complex and involves multiple molecular pathways characterized by numerous genetic and epigenetic lesions^5^. CRC can arise from both hereditary and non-hereditary sporadic mutations, but more than 85% of CRC are non-familial. The inactivation of the adenomatous polyposis coli (APC) or its downstream signaling components are common in both hereditary and non-hereditary CRCs. These are also among the best-understood pathways involved in the initiation of CRC^4^. However, other genetic and epigenetic alterations are required for the full development of CRC^5^. Loss of APC function results in the accumulation of β-catenin, resulting in the transcription of a large number of cancer causing target genes^6, 7^, alteration of cell-cell adhesion and cell migration^8^. Despite these noticeable functions of APC, its functional loss alone in both mouse and human is not sufficient to account for the full blown development of CRC^9^, suggesting a significant involvement of other pathways in the tumorigenesis of CRC.

Transmembrane and immunoglobulin domain containing (TMIGD) family proteins are newly identified class of immunoglobulin (Ig) domain containing cell adhesion molecules (Ig-CAMs), which include TMIGD1, immunoglobulin and proline rich receptor-1 (IGPR-1)^10-12^ and TMIGD3^13^, which also acts as a tumor suppressor^14^. Although originally described as a protector of renal epithelial cells from oxidative cell injury^15^, TMIGD1 is downregulated in human renal cancer and its re-expression in renal tumor cells inhibits cell proliferation, migration and tumor growth in mouse tumor xenograft^15^. More importantly, a recent whole genome sequencing of pre-invasive colorectal tumor revealed that expression of TMIGD1 is progressively lost in colon cancer^16^. While the change of TMIGD1 expression in colon cancer has been documented, the functional importance of loss of TMIGD1 in human colon cancer remains to be elucidated. In this study, we demonstrate that the loss of TMIGD1 in mice results in the development of intestinal adenomas. We provide evidence that TMIGD1 acts as a tumor suppressor by arresting cell cycle at G2/M. Our findings provide novel insights into the pathogenesis of colorectal cancer and a possible therapeutic target.

## Materials and Methods

### Antibodies, Plasmids and Primers

Anti-TMIGD1 antibody was previously described ^15^. The following antibodies was also used in this study. PCNA antibody (cat# ab2426) was purchased from abcam. ZO1 antibody (cat# 339100) was purchased from life technologies. Actin (cat# 4968S), β-Catenin (cat# 8480P), CDX2 (cat# 3977S), and E-Cadherin (cat# 14472S) antibodies were purchased from Cell Signaling Technologies. Villin antibody (cat# sc-58897) was purchased from Santa Cruz Biotechnologies. TMIGD1 was cloned into retroviral vector, pMSCV-puro and described before (Arafa et al., 2015).

### Cell culture

HCT116, DLD1, HT29 and RKO cells were grown in RPMI medium plus 10% FBS and penicillin/streptomycin. Cells were purchased from ATCC.

### Mouse studies

TMIGD1 heterozygous mice was generated in Nanjing BioMedical Research Institute of Nanjing University (NBRI), China, which is in the background of C57BL/6J mouse. With subsequent breeding, we generated homozygous TMIGD1 mice. All mice used in this study were bred and maintained at Boston University Medical Center after approval from Institutional Animal Care and Use Committee. The following primers were used for genotyping of TMIGD1 mice. Forward primer: CCCTATATCCTCAGGCTCTG. Reverse primer: CGTTCAGCACTACTGTAACGGAC.

### Preparation of mouse intestine and histology analysis

Mice were euthanized, and the intestinal tissues were harvested. A gavage needle was used to flush the colon and small intestine with ice cold PBS. Organs were then stretched across filter paper and opened longitudinally to fix in 10% formalin overnight at 4°C. Tissues were Swiss rolled with the distal end of the intestine closest to the center of the coil, and the proximal end at the outside. The Swiss rolled intestines were paraffin embedded and sectioned at 4μm for histological examination. Three sections from different segments of the block were stained with H&E and grade by a veterinary pathologist and a surgical pathologist in a blind manner for detection of atypical hyperplasia, adenoma, or adenocarcinoma. Additional sections were used for immunohistochemistry or immunofluorescent staining with antibodies as described in the figure legends.

### Cell cycle analysis

RKO cells expressing empty vector and TMIGD1 were plated onto 10cm tissue culture plates at 70-80% confluence. Cells from each cell line were starved for 0 and 72 hours. For harvesting, cells were collected by trypsinization and washed with PBS. Cells were then fixed with 70% ethanol and stored at 4°C for at least 30 min. After fixation, cells were washed twice with PBS and 1×10^6^ cells per group were re-suspended in PBS supplemented with 50ul of RNase (100ug/ml stock). 10 min before flow cytometry analysis is performed, propidium iodide (5µl/group, 1mg/ml stock) is added to the samples and briefly vortexed. Flow cytometry was performed by BD LSRII and analyzed with FlowJo.

### Immuno-paired-antibody detection (ActivSignal assay) analysis

ActivSignal assay examines phosphorylation or expression of 70 different human protein targets, which covers 20 major signaling pathways (Meyer et al., 2008). RKO cells expressing empty vector (EV) or TMIGD1 were plated in 96-well plates in triplicates and subjected to ActiveSignal Assay analysis as described in the figure legend.

### Tail vein metastasis assay

To evaluate the capacity of CRC cell line, HCT116 cell expressing GFP alone or expressing TMIGD1/GFP to extravasate and grow, equal number of cells (5×10^5^ cells per/mouse) were injected via the tail vein and metastasis to lung was evaluated after two weeks. Specifically, mice were sacrificed, lungs were removed, and slides of frozen tissues were prepared and viewed under a fluorescence microscope.

### Immunofluorescence Microscopy

Mice intestinal tissues were sectioned to 4 microns and baked for 1 hour. Tissue slides were then deparaffinized using decreasing alcohol gradients. Slides were then submerged into sodium citrate buffer for antigen unmasking. Tissue sectioned were then permeabilized using 0.3% Triton X-100. Slides were blocked using 5% BSA in TBST. Primary antibody was then added to tissue sections and were incubated. Fluorescent antibody was added to the tissue sections and was incubated in the dark. Slides were then mounted using VectaShield Antifade Mounting Medium with DAPI.

### Immunohistochemistry (IHC)

Two human colorectal cancer tissue microarray slides (US Biomax, catalog numbers BC05012a and BC05118a) consist of 172 tissue samples (72 on BC05012a and 100 on BC05118a) were stained with anti-TMIGD1 antibody. The patient age, tumor grade, tumor stage and TNM status were provided for each sample on both microarrays. Six samples were excluded from the final analysis due to either poor tissue quality secondary to artifact or complete absence of CRC tissue in the specimen. IHC was performed on both microarray slides using validate TMIGD1 antibody ^15, 17^ and Polymer-HRP secondary antibody (Abcam). The staining intensity of each specimen was rated independently by two pathologists. Each specimen was rated from 0-3, with “0” representing no TMIGD1 staining and “3” representing the highest intensity of TMIGD1 staining. The staining pattern was described for each specimen as granular, perinuclear or diffuse.

### Statistical Analysis

The mean TMIGD1 staining intensity was stratified based on patient age, tumor grade, tumor stage and T rating in the TNM staging system. The mean intensity values of each sub-group were compared for statistical significance via the ANOVA with Tukey post-hoc test. The alpha value for significance was set at p < 0.05.

## RESULTS

### Loss of TMIGD1 in mice causes intestinal adenoma

We first examined expression of TMIGD1 in normal human and mouse intestinal tissues via immunofluorescence staining using a previously validated anti-TMIGD1 antibody ^15, 17^. TMIGD1 was detected at the membranous regions of intestinal epithelial cells in human (**S. Figure 1B, 1C**). Consistent with its previously described characteristic as a cell adhesion molecule, TMIGD1 co-localizes with E-cadherin in human intestinal epithelial cells, a well-characterized marker of epithelial cell adherens junctions (**S. Figure 1D-F**), but not with the tight junction protein, ZO-1 (**S. Figure 2A-C**). Similar to its expression in human intestinal tissue, TMIGD1 also is expressed in mouse intestinal epithelial cells (**S. Figure 3B**).

We next sought to examine overall TMIGD1 expression in human intestinal tissue via genevestigator dataset (https://genevestigator.com). Our analysis revealed that TMIGD1 mRNA is highly expressed in human colonic tissues, including cecum, small intestine, large intestine and jejunum (**S. Figure 4A**). Additionally, analysis of the RNA sequence of 19 human fetus tissues (NIH Roadmap Epigenomics Mapping Consortium via (https://www.ebi.ac.uk/gxa/home) similarly demonstrated that TMIGD1 mRNA level is highest in the intestinal tissues followed by the kidney (**S. Figure 4B**). The TMIGD1 mRNA in other organs, including heart, stomach, muscle, adrenal gland and placenta were either not expressed or expressed at very low levels (**S. Figure 4B**).

To investigate the potential function of TMIGD1 in the intestinal function *in vivo*, we generated homozygous TMIGD1 knockout mice via CRISPR/Cas9 system (**Figure 1A**). CRISPR/Cas9-mediated loss of TMIGD1 was confirmed by qPCR (**Figure 1B**). Although, TMIGD1-/- mice are viable, fertile and pups display no apparent abnormality, however, as TMIGD1-/- mice get older (4 months and older), they developed intestinal hyperplasia (**Figure 1C**). Analysis of intestines of TMIGD1-/- mice revealed that 80% (8 out 10) of TMIGD1-/- mice develop polyps in small intestine, large intestine and rectum (**Figure 1C**). The average number of polyps observed were between 7-12/mice. Furthermore, H&E analysis demonstrated the development of microtubular adenoma and tubular adenoma in TMIGD1-/- mice (**Figure 1D**).

**Figure 1.**
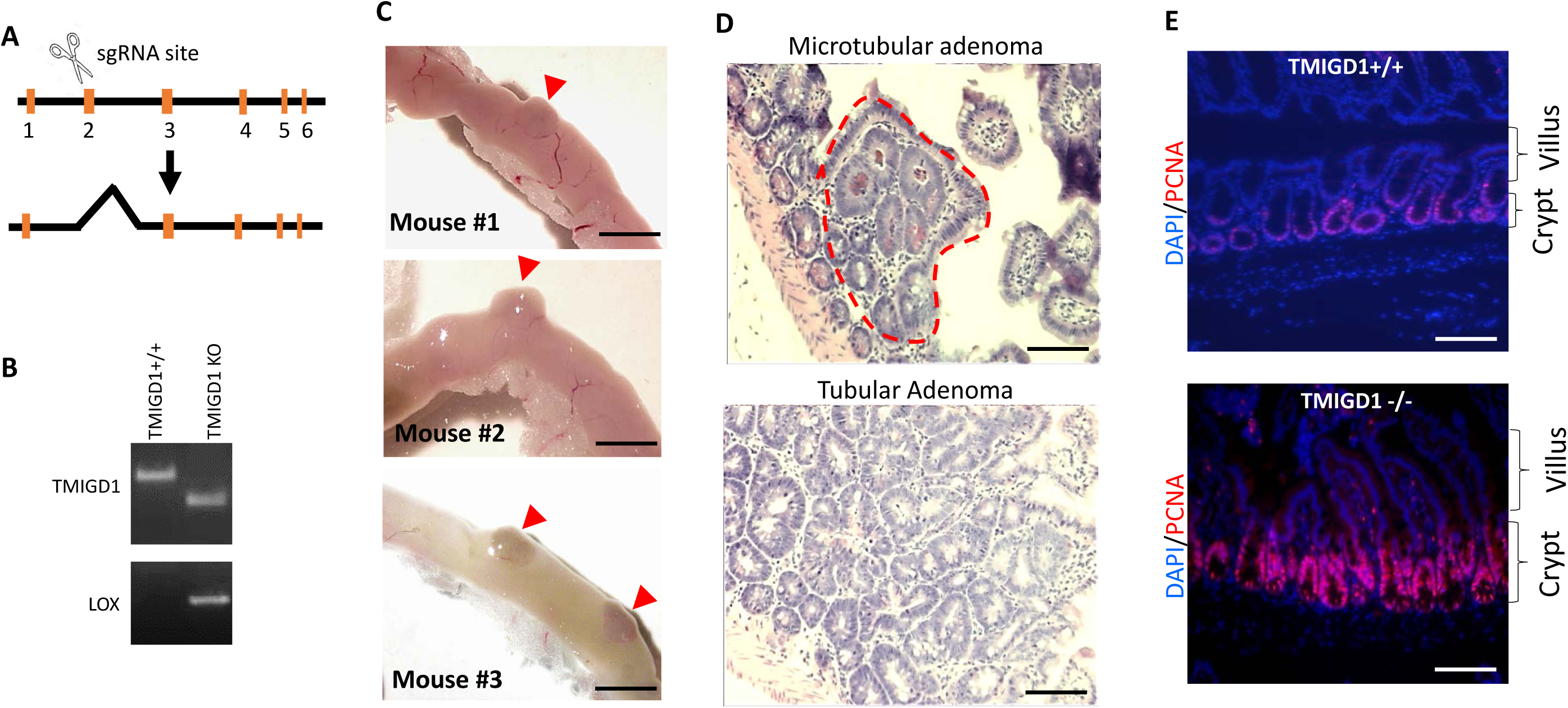
Loss of TMIGD1 in mice causes intestinal adenoma. (**A**) Schematic of sgRNA-mediated knockout of TMIGD1. (**B**) qPCR analysis of wild type and TMIGD1 KO mice. (**C**) Representative images of formation of intestinal adenoma in TMIGD1 KO mice. (**D**) H&E staining of intestinal tissue of TMIGD1 KO. (**E**) Immunofluorescence staining of wild type and TMIGD1 KO mice intestines with PCNA antibody (red, PCNA, blue, DAPI). Image magnification 50µM.

It is highly likely that development of adenoma in TMIGD1-/- mice is associated with aberrant cell proliferation. Therefore, we examined intestinal tissues of WT and TMIGD1 KO mice with proliferating cell nuclear antigen (PCNA), a marker for cell proliferation. Immunostaining of the intestinal tissues of TMIGD1+/+ and TMIGD1-/- mice for PCNA showed that loss of TMIGD1 in mice results in the hyper-proliferation of the intestinal epithelial cells (**Figure 1E**). While there was weak staining for PCNA in the intestinal epithelium (in the crypt region) of TMIGD1+/+ mice, there was strong staining for PCNA in the intestinal epithelium of TMIGD1-/- mice (**Figure 1E**). Furthermore, Ki67 staining, another marker for *in vivo* cell proliferation, showed similar result (data not shown). TMIGD1-/- mice are currently being monitored for potential adenocarcinoma development and other potential pathologies as they get older. Taken together, we demonstrate that loss of TMIGD1 in mice causes intestinal adenoma, with a significant potential role in human colorectal cancer.

### Loss of TMIGD1 in mice alters apico-basal organization and maturation intestinal epithelial cells

Our examination of intestinal tissues of TMIGD1-/- mice further demonstrated that in addition to development of adenoma, overall organization of the intestinal epithelium in TMIGD1-/- mice was abnormal. Specifically, H&E staining revealed that brush border, an actin-based membrane protrusions known as microvilli, in TMIGD1-/- mice was distinctively reduced. While brush border in the wild type mouse intestine was clearly evident and well organized, the brush border in the intestinal tissues of TMIGD1-/- mouse was mostly absent (**Figure 2A**). To further investigate the apparent loss of brush border in TMIGD1-/- mice, we stained the wild type and TMIGD1 -/- mice intestinal tissues with actin and villin, which are common markers for the brush border. Immunofluorescence staining of wild type intestine with actin and villin demonstrated that both actin and villin are uniformly present at the brush border of wild type mice (**Figure 2B, 2C**). However, in TMIGD1-/- mice actin localization at the brush border was significantly reduced and was aberrantly localized at the crypt and villus regions (**Figure 2B**). Interestingly, villin localization to brush border was significantly reduced or mislocalized in TMIGD1-/- mice (**Figure 2C**). To obtain additional insight about the brush border impairment in the intestinal epithelial cells in TMIGD1 KO mice, we stained the intestinal tissues of wild type and the TMIGD1 KO mice with atypical protein kinase C (aPKC), a marker for apico-basal polarity. The data demonstrated that aPKC is uniformly present at the brush border in the wild type mice but not in the TMIGD1 KO mice (**Figure 2D**).

**Figure 2.**
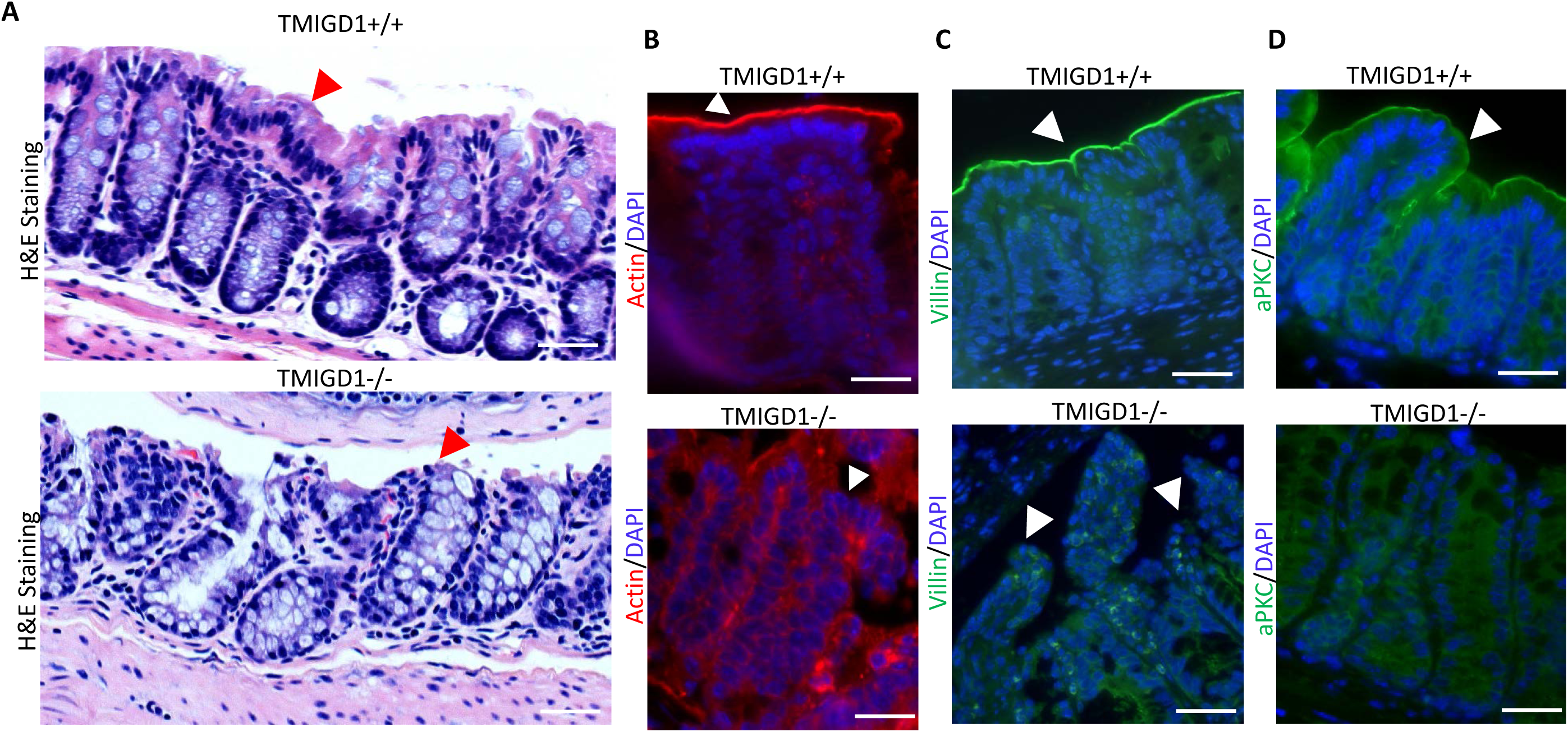
Loss of TMIGD1 in mice impairs brush border development. (**A**) H&E staining of wild type and TMIGD1 mice. Arrows (red) point to brush border region of intestine from wild type and TMIGD1 KO mice. (**B**) Immunofluorescence staining of intestine from wild type and TMIGD1 KO mice with β-actin antibody (red, actin, blue, DAPI). Arrows (white) point to brush border regions in wild type and TMIGD1 KO mice. (**C**) Immunofluorescence staining of intestine from wild type and TMIGD1 KO mice with Villin antibody (green, Villin, blue, DAPI). Arrows (white) point to brush border regions in wild type and TMIGD1 KO mice. (**D**) Immunofluorescence staining of intestine from wild type and TMIGD1 KO mice with aPKC antibody (green, blue, DAPI). Image magnification (A-D) 50µM.

Furthermore, staining with the E-cadherin antibody demonstrated that in the wild type mouse, E-cadherin was strongly positive both at the crypt and microvilli of epithelium (**Figure 3A**), which indicates a normal cellular junctions of epithelium. Similarly, β-catenin staining displayed a similar pattern (**Figure 3B**). However, although the crypt epithelial cells were positive for E-cadherin in TMIGD1-/- mice, it was diffusely present at the cell junctions and instead it was prominently detected at the apical membrane (**Figure 3A**) and was mostly absent in the brush border (**Figure 3A**). Moreover, β-catenin was diffusely present at the intestinal cell junctions in TMIGD1-/- mouse (**Figure 3B, lower panel**). E-cadherin is known to localize to the lateral membrane of differentiated epithelial cells, providing the structural foundation for adherens junctions, which also promotes epithelial apical–basal polarization ^18^. E-cadherin also is required for maturation of intestinal Paneth cells ^19^, and β-catenin is thought to play a central role in maturation and morphogenesis of crypt and villus ^20, 21^. Additional staining with zona occluden protein (ZO1), a marker for tight junction that is also required tight junction formation of epithelial cells ^22^, demonstrated that in TMIGD1-/- mouse, ZO1 is largely absent at the junctions of intestinal epithelial cells (**Figure 3C, upper panel compared to lower panel**).

**Figure 3.**
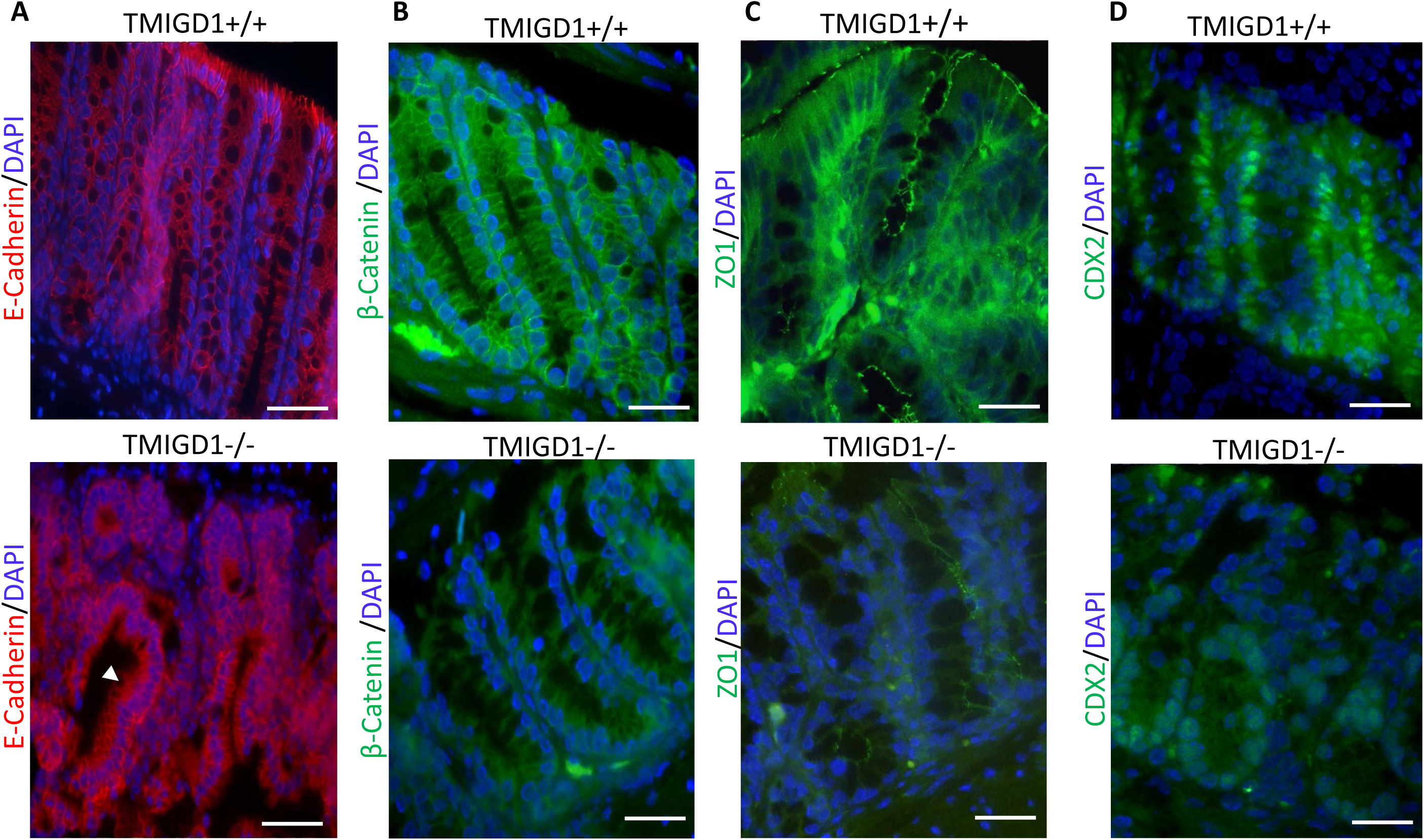
Loss of TMIGD1 alters intestinal epithelium organization and maturation. (**A**) Immunofluorescence staining of intestines of wild type and TMIGD1 KO mice with E-cadherin antibody (red, E-cadherin, blue, DAPI). Arrow (white) points to mislocalization of E-cadherin in the intestinal epithelium in TMIGD1 KO mouse. (**B**) Immunofluorescence staining of intestines of wild type and TMIGD1 KO mice with E-cadherin-associated protein β-catenin antibody (green, β-catenin, blue, DAPI). (**C**) Immunofluorescence staining of intestines of wild type and TMIGD1 KO mice with ZO1 antibody, a tight junction marker protein (green, ZO1, blue, DAPI). (**D**) Immunofluorescence staining of intestines of wild type and TMIGD1 KO mice with CDX2 antibody, a marker for intestinal epithelium marker(green, CDX2, blue, DAPI). Image magnification (A-D) 50µM.

To gain further insights into a possible mechanism for aberrant junctional development of intestinal epithelium in TMIGD1 KO mice, we questioned whether loss of TMIGD1 in mice alters the intestinal epithelial cell maturation. To this end, we stained the TMIGD1-/- mouse intestinal tissue with CDX2, a marker for maturation of intestinal epithelial cells including goblet and Paneth cells ^23-25^. While, the CDX2-positive epithelial cells were uniformly present at the crypt and villus of the wild type mice, in TMIGD1-/- mouse, CDX2 positive cells were detected only at the lower part of the crypts, albeit, weakly and inconsistently (**Figure 3D, upper panel compared to lower panel**). Altogether, the data demonstrate that the loss of TMIGD1 in mouse impairs intestinal epithelial cell maturation and intercellular junctions.

### TMIGD1 arrests the cell cycle at G2/M phase and inhibits cell migration and metastasis

Considering that loss of TMIGD1 in mice induces intestinal adenomas and increased cell proliferation (**Figure 1**), we decided to investigate the effect of ectopic expression of TMIGD1 in colon cancer cells. To this end, we retrovirally expressed TMIGD1 or empty vector (EV) in RKO cells. Expression of TMIGD1 in RKO cells was confirmed by Western blot analysis using anti-TMIGD1 antibody (**Figure 4A**). We initially measured the effect of expression of TMIGD1 in RKO cells in cell proliferation. Over-expression of TMIGD1 in RKO cells significantly inhibited cell proliferation (**Figure 4B**). While proliferation of RKO cells expressing empty vector (EV/RKO) increased in a time dependent manner, proliferation of RKO cells expressing TMIGD1 (TMIGD1/RKO) was reduced by 53% at day 4 (p=0.0033, number of cell for EV/RKO cells was 66.5×10^4^ vs. 30.8×10^4^ for TMIGD1/RKO cells) (**Figure 4B**). We postulated that TMIGD1 could inhibit cell proliferation by a mechanism that modulates the cell cycle. Therefore, we examined the cell cycle distribution of EV/RKO and TMIGD1/RKO cells by FACS (LSRII) analysis via propidium iodide staining. TMIGD1 expressing RKO cells showed a significant cell cycle arrest at G2/M phase (25.4% versus 9.3%) of the cell cycle in the presence of 10% FBS (**Figure 4, Panel C compared to panel D**) and serum-starved (72hr) 14.1% versus 7.2% (**Figure 4, panel E compared to panel F**). Accumulation of G2/M phase of the cell cycle of TMIGD1/RKO cells followed by a markedly decreased G0/G1 phase of the cell cycle (**Figure 4, Panel C compared to panel D and panel E compared to panel F**).

**Figure 4.**
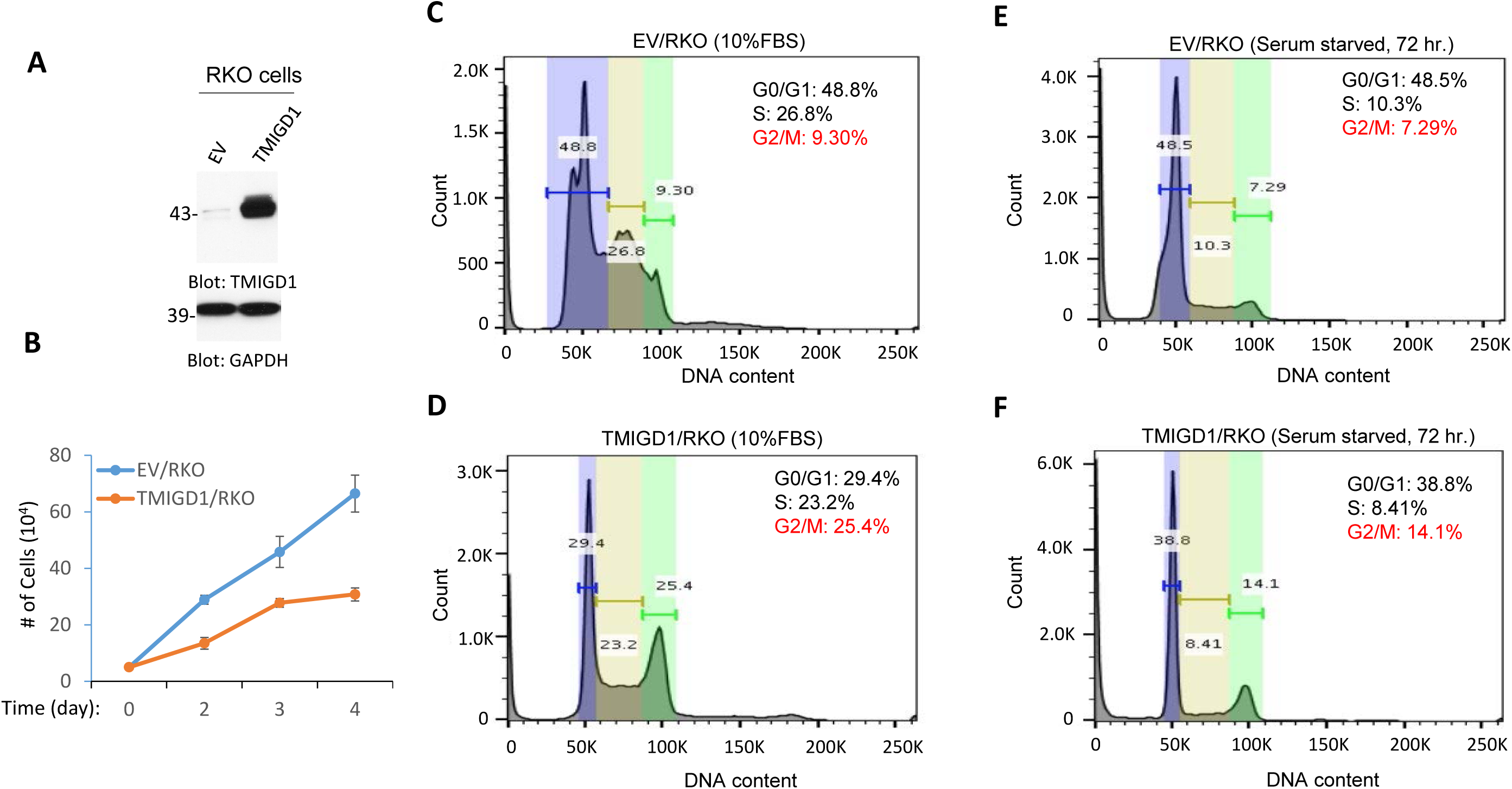
Re-expression of TMIGD1 in RKO cells inhibits cell proliferation and results in cell cycle arrest at G2/M phase. (**A**) Western blot analysis of TMIGD1 expression in RKO cells. (**B**) Proliferation of RKO cells expressing empty vector (EV) or TMIGD1 was determined by direct counting of cells by hemocytometer. (**C—F**) Equal number of RKO cells expressing empty vector (EV) and TMIGD1 about 70-80% confluence were kept in 10%FBS or starved for 72 hours. Cells were fixed with 70% ethanol, stained with propidium iodide and analyzed by BD LSRII and graphs were generated with FlowJo.

To investigate the potential mechanism by which TMIGD1 affects cell cycle in RKO cells, we analyzed RKO cells expressing TMIGD1 for activation of twenty major cancer pathways consisting of 70 individual proteins via a recently developed immuno-paired-antibody detection system (ActiveSignal Assay) analysis platform (Meyer et al., 2018). Among the major pathways that were affected by TMIGD1 in RKO cells were the proteins that are known to inhibit cell cycle and cell proliferation. Specifically, TMIGD1 upregulated expressions of p21CIP1 (cyclin-dependent kinase inhibitor 1) and p27KIP1 (cyclin-dependent kinase inhibitor 1B expression) (**Figure 5A**), proteins whose expression are critically important for negative regulation of the cell cycle progression (Abukhdeir and Park, 2008). Furthermore, TMIGD1 reduced phosphorylation of CyclinD1 and Cyclin-dependent kinase 1 (CDK1/Cdc2), increased phosphorylation of retinoblastoma protein (Rb), and p38MAPK (MAPK14) (**Figure 5A**). Western blot analysis further confirmed the effect of TMIGD1 over-expression on the phosphorylation of p38, phosphorylation of Rb and induction of p21 and p27 (**Figure 5B**). Taken together, the data suggests that TMIGD1 expression in RKO cells induces growth suppression via cell cycle arrest at G2/M phase.

**Figure 5.**
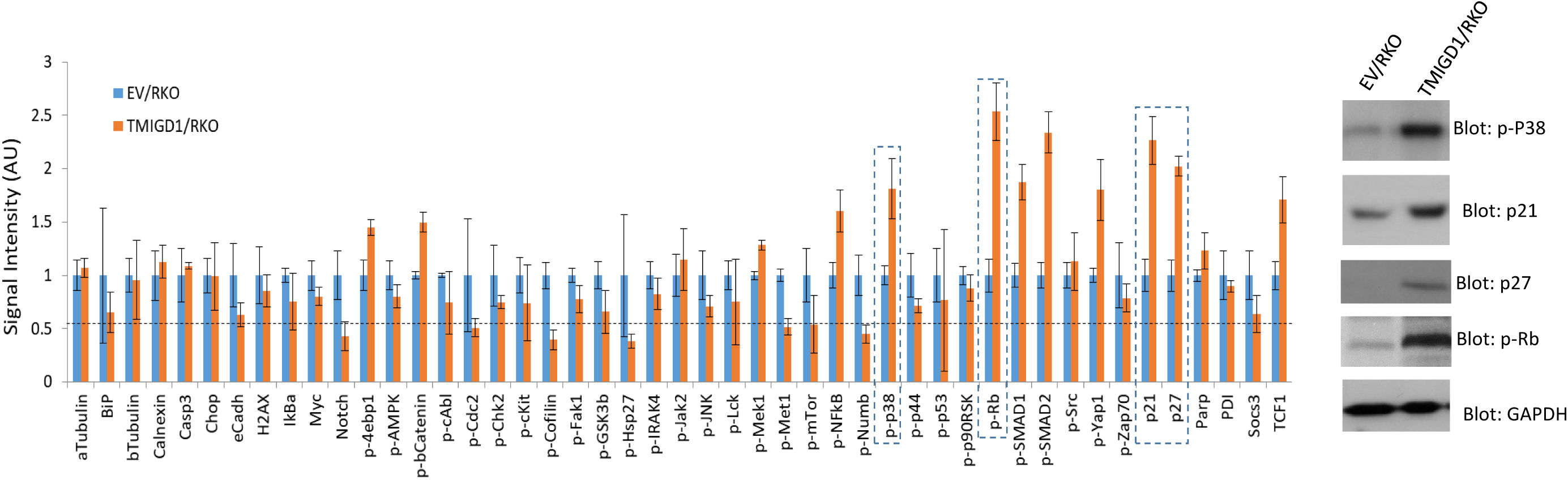
Re-expression of TMIGD1 in CRC cells inhibits cell migration and metastasis. (**A & B**) Migration of RKO and HCT116 cells expressing empty vector or TMIGD1 was determined in Boyden chamber coated with Matrigel. (**C**) Tail vein mouse metastasis assay of HCT116 cells expressing GFP or TMIGD1/GFP (4 mice/group). Metastasis of cells to mouse lung is shown. Image J software was used to quantify GFP-positive cells in mouse lung. (**D**) Western blot analysis of TMIGD1 expression in HCT116 cells.

To examine the effect of TMIGD1 in tumorigenesis beyond cell proliferation and cell cycle, we measured the effect of TMIGD1 in cell migration in RKO and HCT116 cells. Expression of TMIGD1 in RKO and HCT116 cells inhibited cell migration (**Figure 6A, B**). Next, we examined the role of TMIGD1 in inhibition of tumor cell invasion in mouse. For this purpose, we generated GFP/HCT116 cells expressing TMIGD1 or empty vector and assessed their metastatic potentials in the experimental metastasis mouse model via tail vein injection. Mice injected with EV/GFP/HCT116 cells showed significant (5 out 5) metastasis to lung as determined by the presence of GFP positive cells in lung. However, the metastasis of TMIGD1/GFP/HCT116 cells (P= 0.0034) was significantly less (**Figure 6C**). Expression of TMIGD1 in HCT116 cells also shown (**Figure6D**). The data demonstrate that in addition to the inhibition of cell proliferation, TMIGD1 expression in CRC tumor cells also inhibits tumor migration and metastasis.

**Figure 6.**
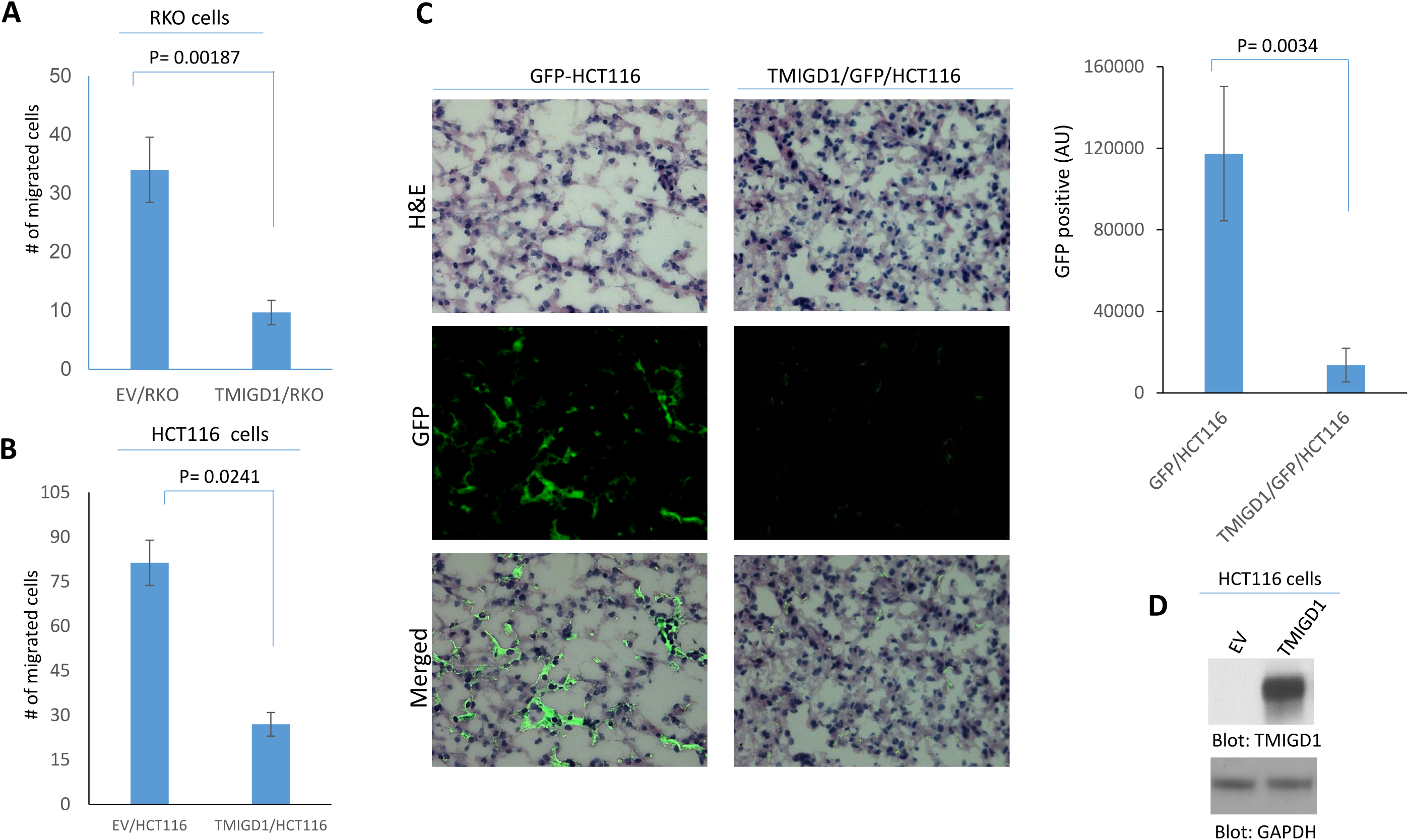
TMIGD1 is downregulated in advanced human colon cancer. (**A**) Two human colorectal cancer tissue microarray slides consist of 172 tissue samples were stained with anti-TMIGD1 antibody. Staining was scored as 0 (negative, <5% cells positive), 1+ (6-25% cells positive), 2+ (26-50% cells positive), and 3+ (>50% cells positive). The tumor differentiation was graded morphologically (grade 1, 2 and 3 as well, moderately and poorly differentiated). (**B**) The average staining intensities were compared using ANOVA with Tukey post-hoc test per tumor grade. (**C**) The average staining intensities were compared using ANOVA with Tukey post-hoc test per tumor differentiation. (**D**) Western blot analysis of TMIGD1 expression in human renal and colorectal cancer cell lines. (**E**) qPCR analysis of TMIGD1 expression in CRC cell lines (HT29, HCT116 and RKO). Human kidney epithelial cells, HK2 was used as a positive control.

### TMIGD1 is downregulated in colon cancer and its downregulation correlates with poor survival

Considering that the loss of TMIGD1 in mice resulted in the development of adenoma, altered epithelial phenotype, and increased cell proliferation, we asked whether TMIGD1 expression is altered in human cancers. Our initial analysis of TMIGD1 mRNA across 23 major human cancer types from The Cancer Genome Atlas (TCGA) via TEMER online site (https://cistrome.shinyapps.io/timer/) ^26^ demonstrated that TMIGD1 mRNA is significantly reduced in human colon, rectum and renal cancers compared to their corresponding normal tissues (**S. Figure 5A**). TMIGD1 expression in other tissues, both normal and cancer, was significantly low (**S. Figure 5A**). Consistent with the downregulation of TMIGD1 in human colon cancer, analysis of 46 human colon cancer cell lines via The Cancer Cell Line Encyclopedia, CCLE ^27^ also indicated that TMIGD1 mRNA was downregulated in colon cancer cell lines (data not shown).

We next examined TMIGD1 protein expression levels on the human CRC tissue microarray slides (n=166) via IHC analysis and TMIGD1 levels was semi-quantitatively analyzed. The normal adjacent tissue (NAT) (n=52) had a mean TMIGD1 staining score of 1.87, and grade 1 (n=21), grade 2 (n=62) and grade 3 (n=29) tumors had means of 2.24, 1.52 and 1.14, respectively (**Figure 7A, B**). Tumor differentiation was graded morphologically (grade 1 to 3 as well, moderately and poorly differentiated). The level of differentiation of each tumor was also subjectively assessed based on the number of glands, gland architecture and cell morphology; samples were rated along a spectrum of “well-,” “moderately-,” or “poorly-differentiated.” Interestingly, TMIGD1 expression was higher in well-differentiated tumors compared to adjacent normal tissue, but the differences were not statistically significant (**Figure 7A, C**). However, expression of TMIGD1 was progressively downregulated with tumor progression as the lowest level of TMIGD1 was observed in the poorly differentiated tumors (**Figure 7A, B**). There were no significant differences observed when TMIGD1 staining intensity was stratified by patient age or tumor stage (data not shown). Furthermore, TMIGD1 expression in multiple colon cancer cell lines (HT29, HCT116, RKO and DLD1) and renal cancer cell lines (786-0, CAKI-1, A498 and TK10) as determined by Western blot analysis showed that TMIGD1 protein levels both in colon cancer and renal cancer cell lines was either very low or undetectable (**Figure 7D**). However, in two colon cancer cell lines (HCT116 and HT29) a protein band at the lower molecular weight was also detected (**Figure 7D**), which may correspond to partially/unglycosylated TMIGD1 (**Figure 7D**). Additionally, qPCR analysis of TMIGD1 mRNA revealed that TMIGD1 mRNA is expressed at low levels in HT29, HCT116 and RKO cells compared to normal human kidney epithelial (HK2) cells (**Figure 7E**). Downregulation of TMIGD1 in CRC prompted us to question whether TMIGD1 expression profile correlates with CRC patient’s survival. We carried out a Kaplan Meier survival analysis of the publically available CRC TCGA dataset. The result revealed that CRC patients with high TMIGD1 expressing tumors had significantly better median overall survival compared with those with low TMIGD1 expressing tumors (**S. Figure 5B**).

**Figure 7.**
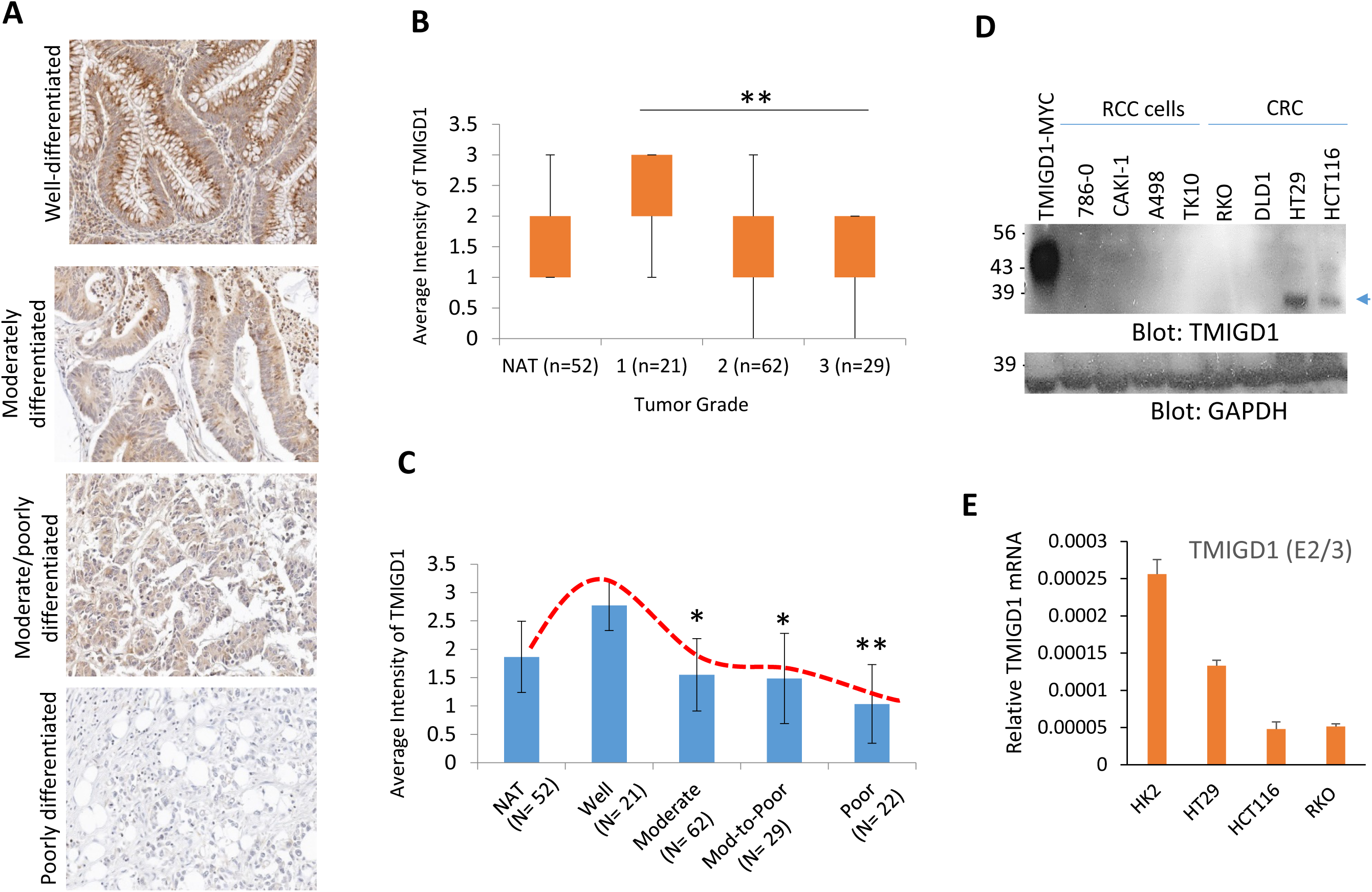

## Discussion

CRC is the second leading cause of cancer morbidity and mortality among men and women. While only about 5-10% of all CRC are genetic, more than 85% of CRC are sporadic. The adenomatous polyposis coli (APC) tumor suppressor gene is mutated in most familial and sporadic colorectal cancers. However, despite its prevalence, mutations in the APC gene are thought to be an early, if not an initiating event, in multistep processes of CRC progression. More importantly, though the mechanism of APC inactivation is fairly well documented, the process by which APC inactivation leads to tumorigenesis remain poorly understood. Our result show that the loss of TMIGD1 in mice increases intestinal epithelial cell proliferation and induces adenoma. We further demonstrated that TMIGD1 is downregulated in human colon cancer and its downregulation is associated with poor survival. Re-expression of TMIGD1 in colon cancer cells inhibited cell proliferation, arrested cell cycle at G2/M phase, and suppressed cell migration and metastasis. Expression of TMIGD1 in CRC cells (RKO cells) induced expression of p^21CIP1^ (cyclin-dependent kinase inhibitor 1), and p^27KIP1^ (cyclin-dependent kinase inhibitor 1B) expression, key cell cycle inhibitor proteins involved in the regulation of the cell cycle, which may account for the effect of TMIGD1 in cell cycle and tumorigenesis. This study uncovers TMIGD1 as a novel tumor suppressor gene and as a potential therapeutic target to treat CRC.

Intestinal epithelium displays strong apical–basal polarity with a highly developed brush border in the apical plasma membrane of absorptive epithelial cells and a similarly highly polarized organization of the other cell types that populate intestine. Another interesting and novel aspect of the current study is the observation that loss of TMIGD1 in mice significantly impaired brush border development and integrity of intestinal epithelium as was evident by almost nonexistence actin and villin at the brush border and altered E-cadherin and ZO1 cellular distribution. This theme was further echoed by our observation that loss of TMIGD1 in mice interfered with the maturation of intestinal epithelium, especially given that the CDX2-positive cells were only sparsely and inconsistently present in the intestinal epithelium.

In summary, we provide several lines of evidences that TMIGD1 is a tumor suppressor gene that plays a central role in intestinal epithelium integrity and proliferation and that loss of TMIGD1 in mice results in hyperplasia. Therefore, strategies to upregulate expression of TMIGD1 in CRC offers a novel therapeutic target.

## Supplemental Data

## ACKNOWLEDGMENTS AND FUNDING

**Funding**: This work was supported in part through grants from the NIH/NCI (R21CA191970, R21CA193958, CTSI grant 1UL1TR001430 (NR) and NCI RO1CA175382, NIH R01 HL132325 and the Boston University Evans Faculty Merit award (VCC). The work was also supported in part by the Malory Fund, Department of Pathology and Laboratory Medicine, Boston University. The authors thank Mostafa Belghasem (Department of Pathology, Boston University) for his help with the histopathology and animal work.

---

**S. Figure 1.**
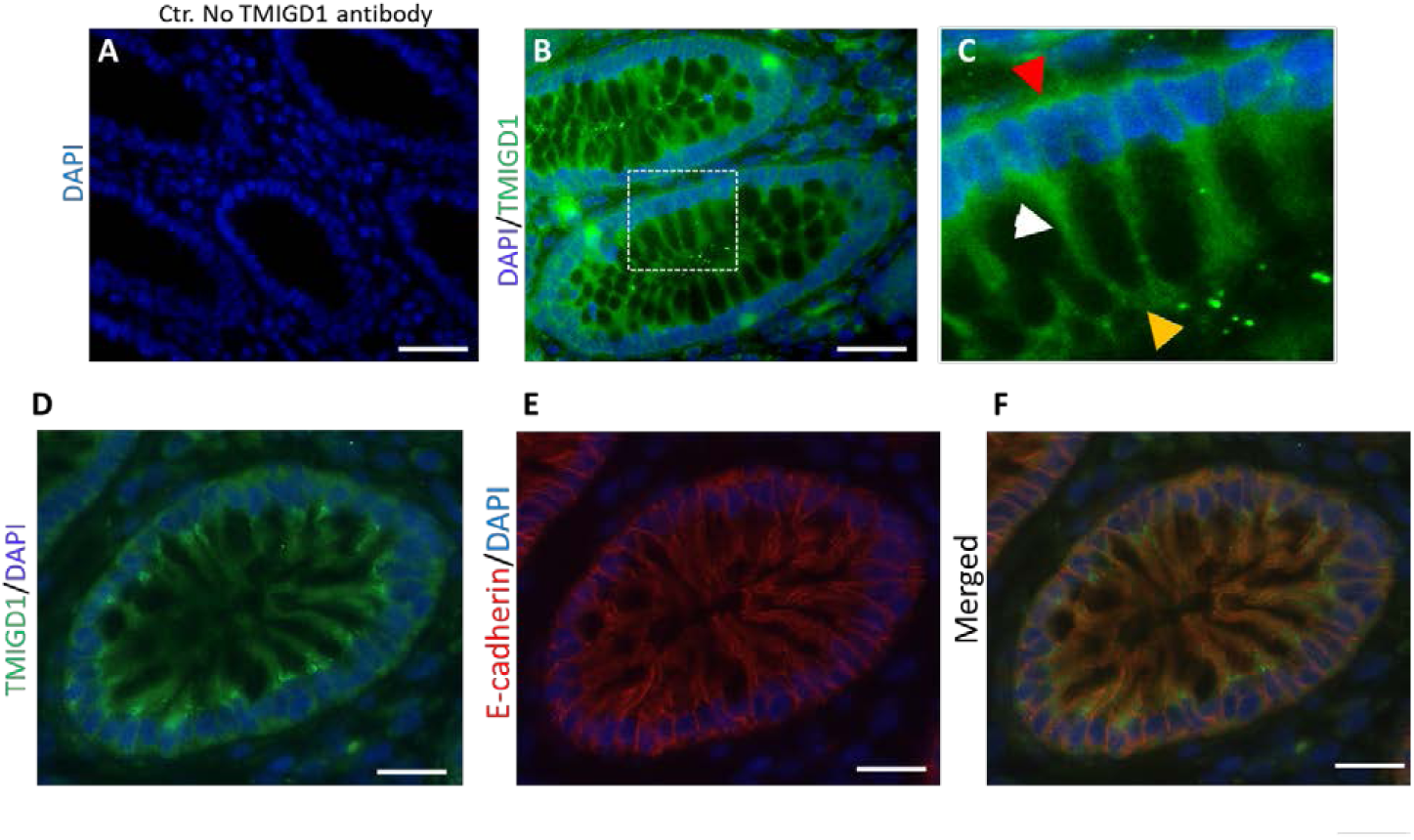
TMIGD1 is expressed in human intestinal epithelial cells and localizes to epithelial cell adherens junctions. (**A**) Normal human intestinal tissue was subjected to immunofluorescence staining without primary antibody. (**B**) The same tissue subjected to immunofluorescence staining using anti-TMIGD1 antibody. (**C**) Enlargement of panel B (dotted white box) is shown. (**A&B**) Image magnification 50µM. (**D-E**) Normal human intestinal tissue was co-stained with anti-TMIGD1 and anti-E-cadherin antibodies. Image magnification 50µM.

**S. Figure 2.**
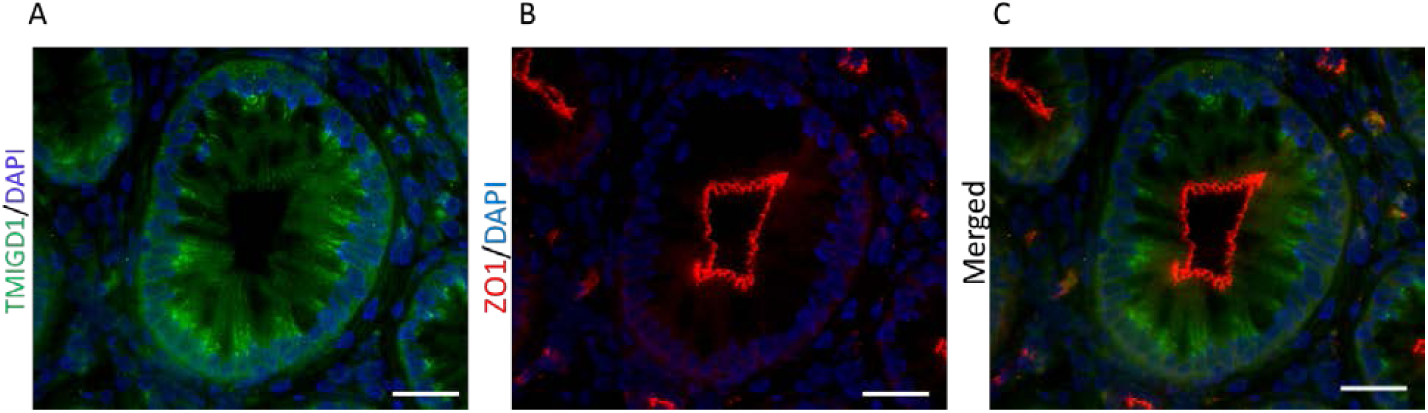
TMIGD1 does not co-localize with the tight junctional protein, ZO1. (**A-B**) Normal human intestinal tissue was co-stained using anti-TMIGD1 antibody and anti-ZO1 antibody. Image magnification 50µM.

**S. Figure 3.**
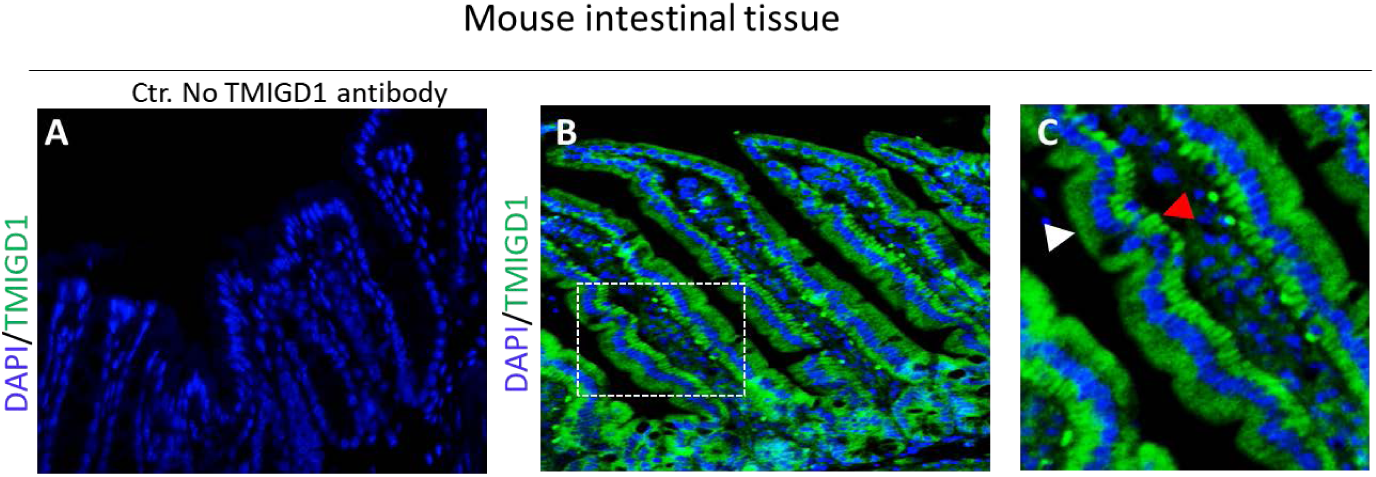
TMIGD1 is expressed in mouse intestinal epithelial cells. (**A**) Normal mouse intestinal tissue was subjected to immunofluorescence staining without primary antibody. (**B**) The same tissue subjected to immunofluorescence staining using anti-TMIGD1 antibody. (**C**) Enlargement of panel B (dotted white box) is shown. White and red arrows indicate basolateral and apical expression of TMIGD1, respectively. Image magnification 50µM.

**S. Figure 4.**
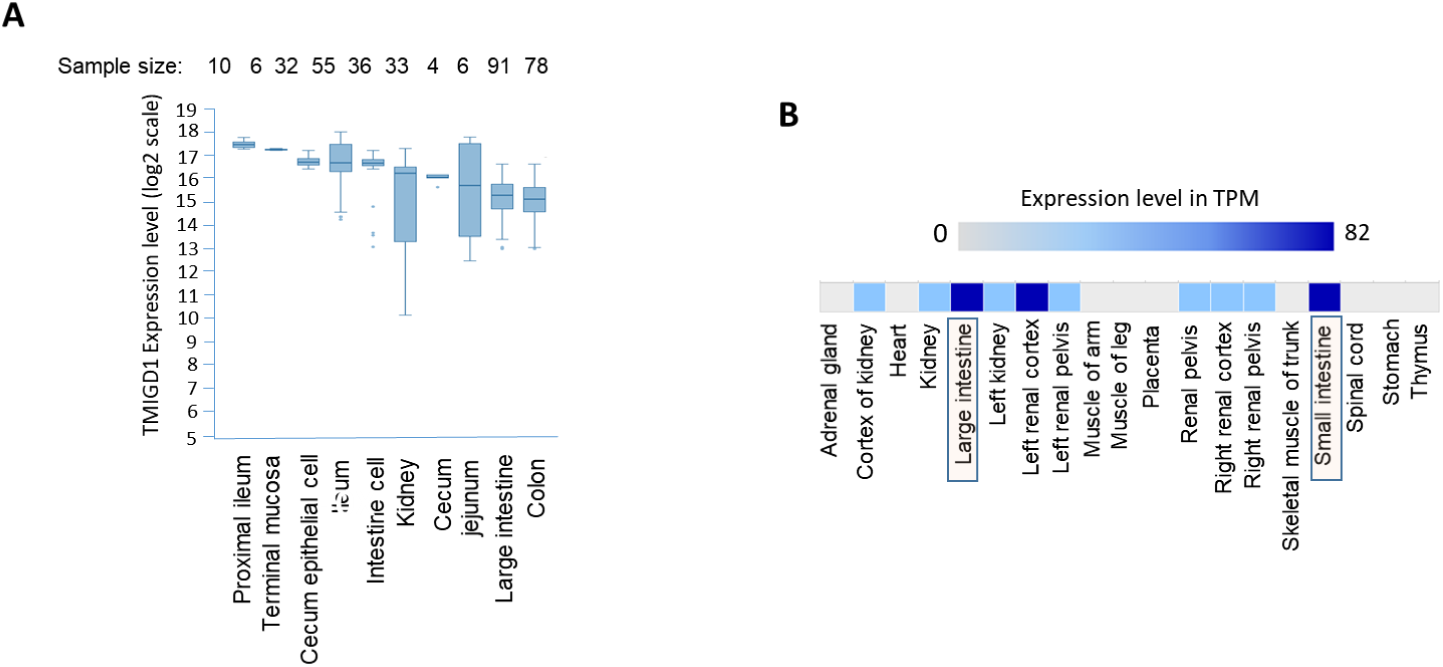
TMIGD1 expression in human and mouse tissues. (**A**) Expression of TMIGD1 in various regions of human intestine. (B) TMIGD1 expression in various mouse tissues/organs.

**S. Figure 5.**
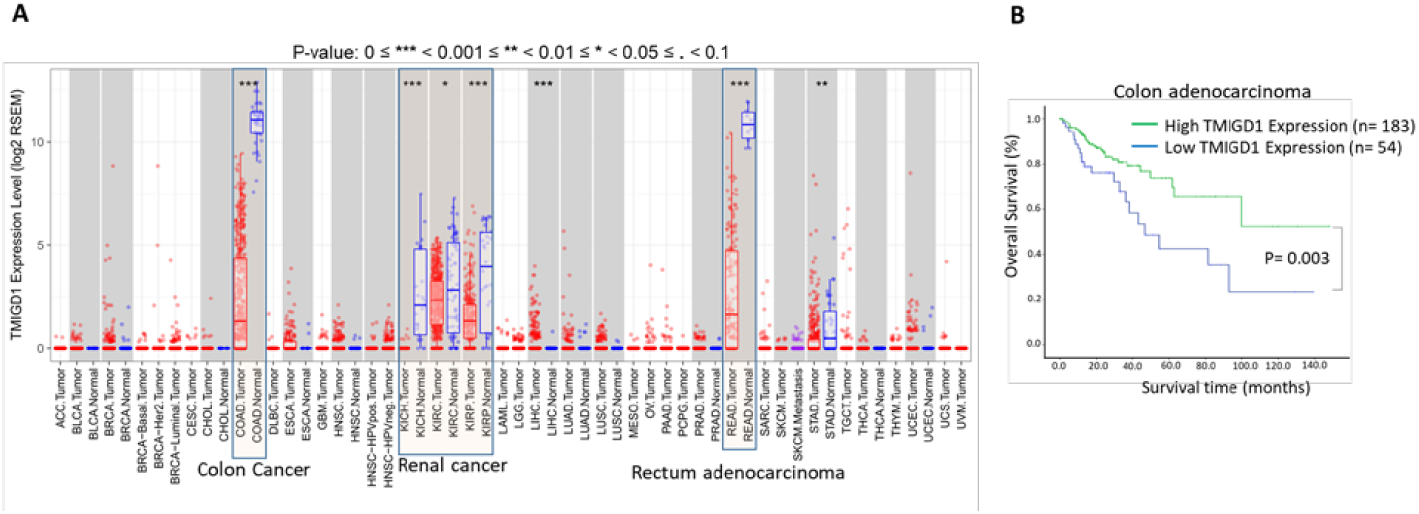
TMIGD1 is downregulated in human colorectal cancer and its downregulation correlates with poor survival. (A) TMIGD1 mRNA levels examined across 23 major human cancer types from The Cancer Genome Atlas (TCGA) via TEMER online site (https://cistrome.shinyapps.io/timer/). (**B**) Kaplan Meier survival analysis via TCGA dataset.

